# Torque generation and regulation in the flagellar motor of *Campylobacter jejuni* differ from those of model organisms

**DOI:** 10.64898/2026.11.10.765344

**Authors:** William H. Hoffmann, Simon Leturcq, Arianna Giannetti, Eli J. Cohen, Morgan Beeby, Francesco Pedaci, Ashley L. Nord

## Abstract

The bacterial flagellar motor is a rotary nanomachine that powers motility by driving the rotation of a helical filament. Most of our mechanistic understanding of this motor derives from studies of model organisms, particularly *Escherichia coli* and *Salmonella enterica* serovar Typhimurium, yet recent structural work has revealed substantial architectural diversity, raising questions about the universality of established biophysical models. Here, we present the first single-motor measurements of torque output and stator dynamics in *Campylobacter jejuni*, a species with a structurally elaborate motor adapted for motility in high-viscosity environments. The *C. jejuni* motor generates an exceptionally high stall torque (∼3600 pN·nm) and, rather than the biphasic torque–speed relationship characteristic of model systems, exhibits an approximately linear decline in torque with increasing speed over the measured range. Time-resolved measurements reveal discrete speed changes, indicating that the stator output remains dynamic despite the presence of a scaffolded stator ring. The inferred number of active stator units averaged 11±4, below the 17-unit complement observed structurally, and showed no clear dependence on viscous load despite substantial motor-to-motor heterogeneity. Following transient collapse of the proton motive force, motor activity recovers rapidly, consistent with reversible activation of pre-associated stator units. Together, these findings reveal a distinct mode of flagellar motor operation, consistent with dynamic regulation of a preassembled stator ring rather than regulation through physical stator exchange, and show that the canonical behavior defined in model organisms is not universal.

## Introduction

The bacterial flagellar motor is a membrane-embedded rotary nanomachine that converts electrochemical ion gradients into torque to drive flagellar rotation and bacterial motility [1, 2]. In-depth analyses of motor mechanical output, including speed, torque, and load-dependent dynamics, have been conducted predominantly in closely related Enterobacteriaceae model organisms *Escherichia coli* and *Salmonella enterica* serovar Typhimurium [1–3]. Yet structural studies over the past decade have revealed extensive diversity among bacterial flagellar motors, establishing the Enterobacteriaceae motor as one lineage-specific architecture among many [4–6]. Despite this diversity, all flagellar motors share conserved stator and rotor components that underlie torque generation: ion flux through membrane-embedded stator complexes drives the cytoplasmic rotor ring (C-ring), and this rotation is transmitted through the axial structure to an extracellular helical flagellar filament that acts as a propeller (Fig. 1a) [1, 2]. The striking structural conservation of these core torque-generating components across divergent bacterial motors suggests that the fundamental mechanism of torque generation is broadly shared [7–10], while leaving open whether the mechanical behavior and regulation of the assembled motor are similarly conserved.

**Figure 1:**
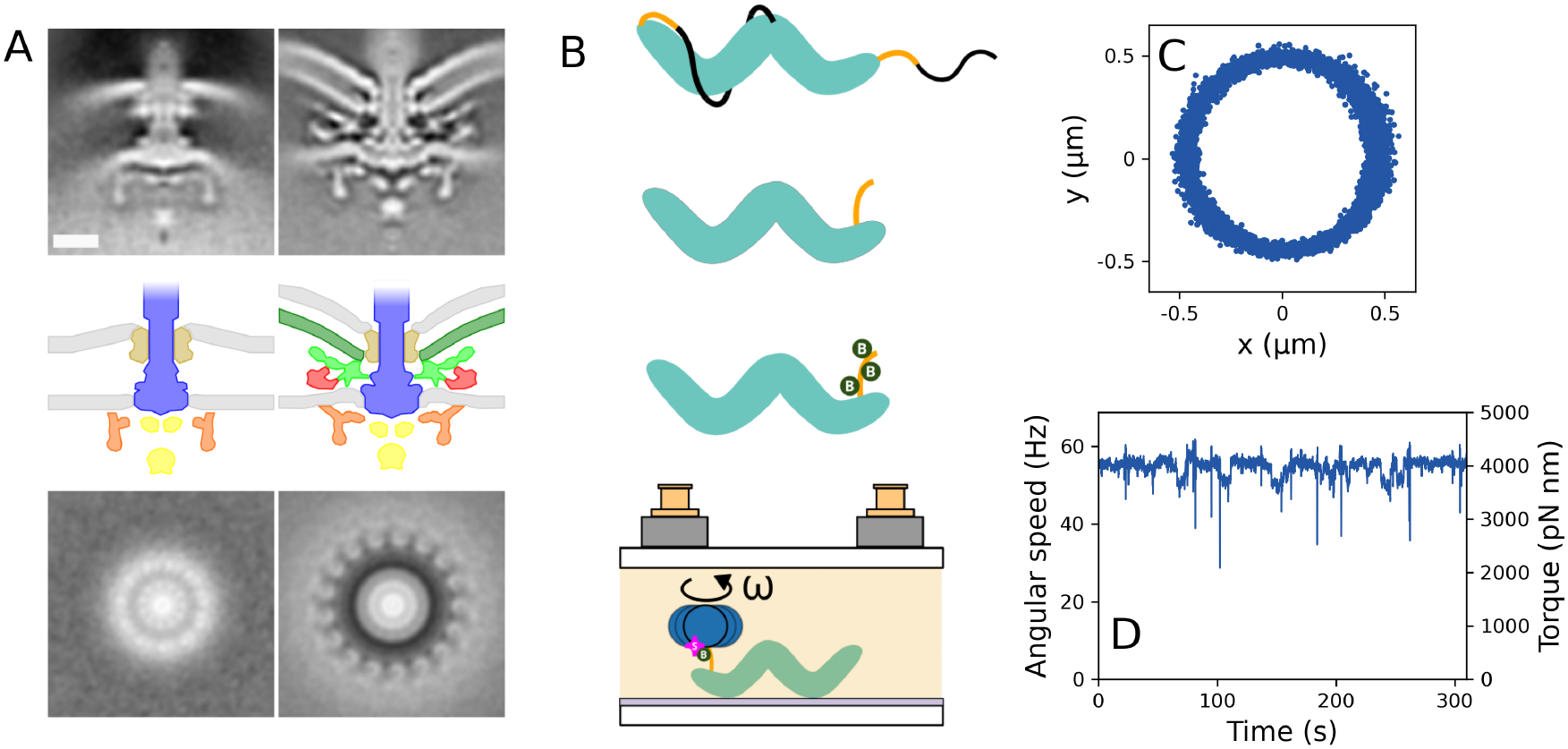
Structure of the motor, bead assay and representative data for the bacterial flagellar motor of *C. jejuni*. A) Comparison of flagellar motor architectures in *S. Typhimurium* (left) and *C. jejuni* (right). Top: side views of flagellar motors obtained by electron cryo-tomography and subtomogram averaging. The scale bar represents 20 nm. Middle: corresponding structural schematics highlighting the conserved motor core and the additional periplasmic scaffold structures in *C. jejuni*. Bottom: axial cross-sections through the motor reconstructions, illustrating the prominent 17-fold arrangement of stator-associated densities in *C. jejuni*, which are not resolved in the *S. Typhimurium* reconstruction. Images and schematics adapted from [4, 5]. B) Schematic of the bead-assay protocol. Cells expressing a shortened, biotin-labelled flagellar filament (green “B” labelled) were attached to a coverslip, and streptavidin-coated beads (magenta “S” labelled) were bound to the filament to report motor rotation. *ω* denotes angular speed. C) Representative *x*–*y* trajectory of a 1400 nm diameter bead attached to a rotating motor. D) Angular speed of the bead trajectory shown in C, with the corresponding torque shown on the right axis. The speed trace was filtered for display as described in the Materials and Methods.

The relationship between torque and rotational speed is a central observable for understanding the internal mechanisms by which rotary molecular motors convert energy into mechanical work. In model organisms, single-motor measurements have revealed a strongly concave-down, biphasic torque-speed relationship, with torque changing little at low speeds before declining more steeply at higher speeds [11–13]. The resulting “knee” in the curve has often been treated as a defining signature of the torque-generating mechanism and is commonly interpreted as a shift from limitation by external viscous load to limitation by intrinsic kinetics of the motor, such as ion translocation or conformational changes [13, 14]. In these organisms, motor torque also adapts to the external load through the dynamic recruitment and release of stator complexes [15–18]. The shape of the torque-speed relationship and load-dependent remodeling of the stator ring have therefore become characteristic features of the current mechanistic framework of flagellar motor operation. However, because nearly all high-resolution measurements of torque-speed behavior and stator dynamics have been performed in these model organisms, it remains unclear to what degree either feature is universal.

The evolutionary history of the Enterobacteriaceae motor further underscores why its behavior may not be representative of flagellar motors more broadly. Phylogenetic analyses suggest that this system was acquired by horizontal gene transfer from an ancestral *β*-proteobacterium and differs from the polar and lateral flagellar systems native to *γ*-proteobacteria [19]. This lineage-specific history motivates comparison with motors that have evolved substantially different architectures. Among these, the *Campylobacter jejuni* motor is one of the most elaborate imaged to date, incorporating multiple periplasmic disk-like structures and an expanded C-ring that interfaces with a scaffolded array of stator units (Fig. 1A) [4, 5, 20]. Cryo-electron tomography reconstructions reveal a 17-fold symmetric stator density [4, 20, 21]. This architecture suggests that, unlike in *E. coli*, stator complexes in *C. jejuni* likely remain associated with the motor rather than dynamically exchange with a membrane pool in response to mechanical load. Yet the mechanical output of the *C. jejuni* motor has never been directly measured at the single-motor level, leaving it unclear whether its distinct architecture is accompanied by differences in torque output, torque-speed behavior, or stator dynamics.

Here, we directly quantify the mechanical output and dynamic behavior of individual *C. jejuni* flagellar motors. We find that the motor generates exceptionally high torque and exhibits a steady, approximately linear decline in torque with increasing speed, rather than the extended constant-torque plateau characteristic of model systems. Time-resolved measurements reveal dynamic changes in stator activity, while rapid recovery following transient collapse of the proton motive force is consistent with reactivation of pre-associated stator units rather than their slower recruitment from a membrane pool. Together, these findings reveal a mode of flagellar motor operation distinct from that described in model organisms and show that canonical torque-speed behavior and regulation through physical stator exchange are not universal features of bacterial flagellar motors.

## Results

### The *C. jejuni* motor generates high torque without an extended constant-torque plateau

We performed bead assay experiments [11, 22] on a genetically engineered *C. jejuni* strain whose motors rotate uni-directionally in the counterclockwise (CCW) direction, allowing us to quantify their torque output and dynamics (Fig. 1B; see Materials and Methods). An example trajectory from a motor measured with a 1400 nm bead is shown in Fig. 1C, with the corresponding angular speed and torque traces shown in Fig. 1D.

Using beads ranging from 250–2700 nm in diameter (measured by transmission electron microscopy; Fig. S1), we quantified motor output over a range of viscous loads spanning more than two orders of magnitude (0.52–78 pN·nm·s/rad). As load increased, the mean rotational speed decreased from 252 ± 78 Hz with 250 nm beads to 7.4 ± 2.3 Hz with 2700 nm beads (Fig. 2A), while the mean torque increased from 770 ± 170 pN·nm to 3520 ± 860 pN·nm (Fig. 2B). Together, these measurements define the population-averaged torque-speed relationship of the *C. jejuni* flagellar motor (Fig. 2C). In contrast to the strongly concave-down canonical CCW torque– speed relationship of *E. coli*, torque in *C. jejuni* decreased monotonically with increasing speed and was well approximated by a linear relation over the measured range. Linear regression yielded an extrapolated stall torque of 3570 ± 200 pN·nm and an extrapolated zero-torque speed of 350 ± 20 Hz. Given the substantial motor-to-motor variability and limited sampling at intermediate loads, we cannot exclude more subtle curvature in the torque-speed relationship; nevertheless, the present data do not support the pronounced concave-down curvature observed in the *E. coli* motor. Individual motor torque-speed measurements illustrating this heterogeneity are shown in the Supplementary Information (Figs. S2 and S3).

**Figure 2:**
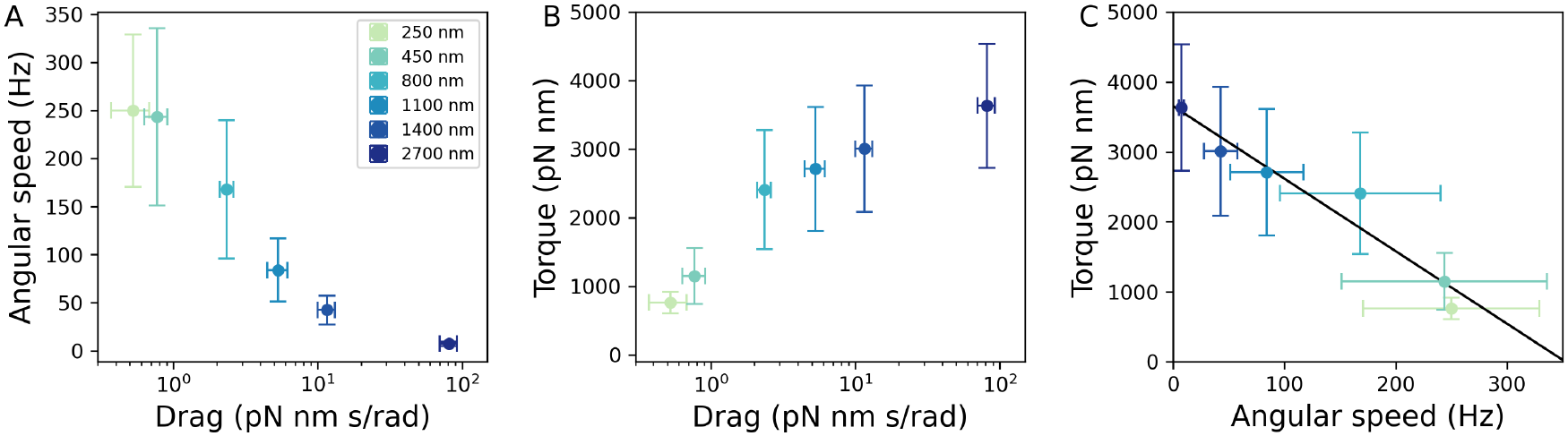
Angular speed and torque output as a function of load on the motor. A) Angular speed versus viscous drag. B) Torque versus viscous drag. C) Torque-speed curve. The black line is a linear regression of torque versus angular speed with a slope of −10.3 ± 1.2 pN·nm/Hz, an extrapolated stall torque of 3570 ± 200 pN·nm, and an extrapolated zero-torque speed of 350 ± 20 Hz. Colored points represent averages for each bead size; corresponding bead diameters are indicated in the legend in A). Error bars indicate standard deviation across motors. Number of motors measured: 27 for 250 nm beads, 42 for 450 nm beads, 13 for 800 nm beads, 29 for 1100 nm beads, 124 for 1400 nm beads, 26 for 2700 nm beads.

### Stepwise speed changes reveal dynamic stator activity across loads

To determine whether the torque-generating activity of the scaffolded stator ring remains dynamic, we examined fluctuations in the time-resolved motor rotation. Across all loads tested, the angular speed traces exhibited frequent and abrupt stepwise changes, qualitatively similar to transitions attributed to stochastic stator association and dissociation in *E. coli* and other species [12, 23–28]. Representative traces obtained with different bead sizes are shown in Fig. 3A–F, with black lines indicating the piecewise-constant levels identified by our step-detection algorithm (see Materials and Methods). The corresponding angular speed histograms (Fig. 3A– F, right) show discrete peaks aligned with the detected levels, supporting their interpretation as distinct rotational states. We quantified the speed-step magnitude, Δ*ω*, across the bead sizes and found that it decreased with increasing viscous drag, *γ* (Fig. 3G), while the corresponding torque increase, Δ*τ* = *γ*Δ*ω*, increased with load (Fig. 3H). Plotting Δ*τ* against Δ*ω* defines a torque–speed relationship derived from individual step events (Fig. 3I), which, under the assumption that each step corresponds to activation or inactivation of a single stator unit, can be interpreted as the torque–speed characteristic of an individual stator unit. Linear regression yielded an extrapolated stall torque of 280 ± 30 pN·nm and an extrapolated zero-torque speed of 42 ± 10 Hz. To estimate the number of functionally active stator units, we calculated the ratio *ω*_0_*/*Δ*ω*, where *ω*_0_ is the absolute motor speed at which each step occurred. The inferred number of active stators showed no clear dependence on viscous load, with bead-size averages ranging from 9 to 14 (Fig. 3J). These values are below the 17 stator-associated positions resolved structurally in the *C. jejuni* motor.

**Figure 3:**
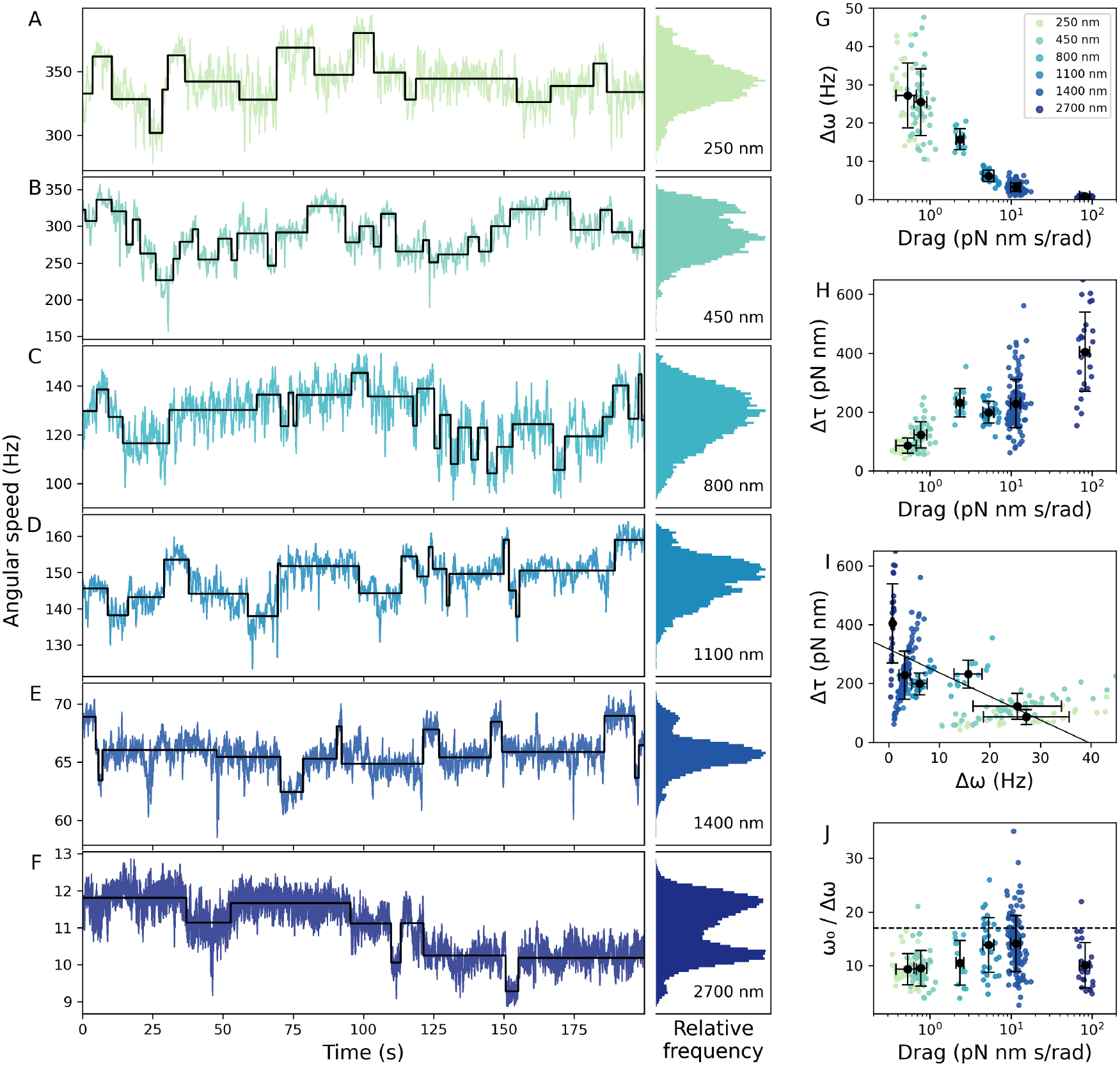
Angular speed and torque dynamics of the flagellar motor of *C. jejuni*. A–F) Angular speed vs. time and corresponding angular speed histograms for individual motors with bead diameters indicated in each histogram plot. Colored lines represent angular speed; black lines show the output of the step-finder algorithm, from which we estimate the number of functionally active stator units (see Materials and Methods). G) Speed step size (Δ*ω*) vs. viscous drag. H) Torque step size (Δ*τ*) vs. viscous drag. I) Δ*τ* vs. Δ*ω*. The black line is a linear regression of the average values. Linear regression yielded a slope of −6.7 ± 1.8 pN·nm/Hz, an extrapolated stall torque of 280 ± 30 pN·nm, and an extrapolated zero-torque speed of 42 ± 10 Hz. J) Ratio of angular speed to step size (*ω*_0_*/*Δ*ω*). For G–J), colored data points represent the average values for individual beads, except *ω*_0_*/*Δ*ω* where the median is plotted, and the black points correspond to the averages at each bead size. Error bars indicate the standard deviation.

### Rapid recovery after PMF restoration is consistent with reactivation of pre-associated stators

To investigate the mechanism underlying the stepwise speed changes described above (Fig. 3), we performed “resurrection” experiments in which the proton motive force (PMF) was transiently dissipated and subsequently restored. In *E. coli*, PMF collapse causes stator units to dissociate from the motor, whereas subsequent PMF restoration leads to their stochastic reassembly, producing stepwise increases in speed and torque [23, 24, 29, 30]. We applied a similar protocol in *C. jejuni* by transiently treating cells with the protonophore CCCP. Rotation halted within seconds of CCCP addition and resumed following washout, ultimately recovering to approximately 85% of the pre-treatment speed.

Recovery kinetics differed markedly between the two species. In *C. jejuni*, speed recovered rapidly, reaching 50% of its pre-treatment value approximately 3 s after the onset of renewed rotation. By contrast, *E. coli* recovered more slowly, reaching the corresponding level after approximately 25 s (Fig. 4A). Representative resurrection traces of *C. jejuni* and *E. coli* motors are shown in Fig. 4B,C.

**Figure 4:**
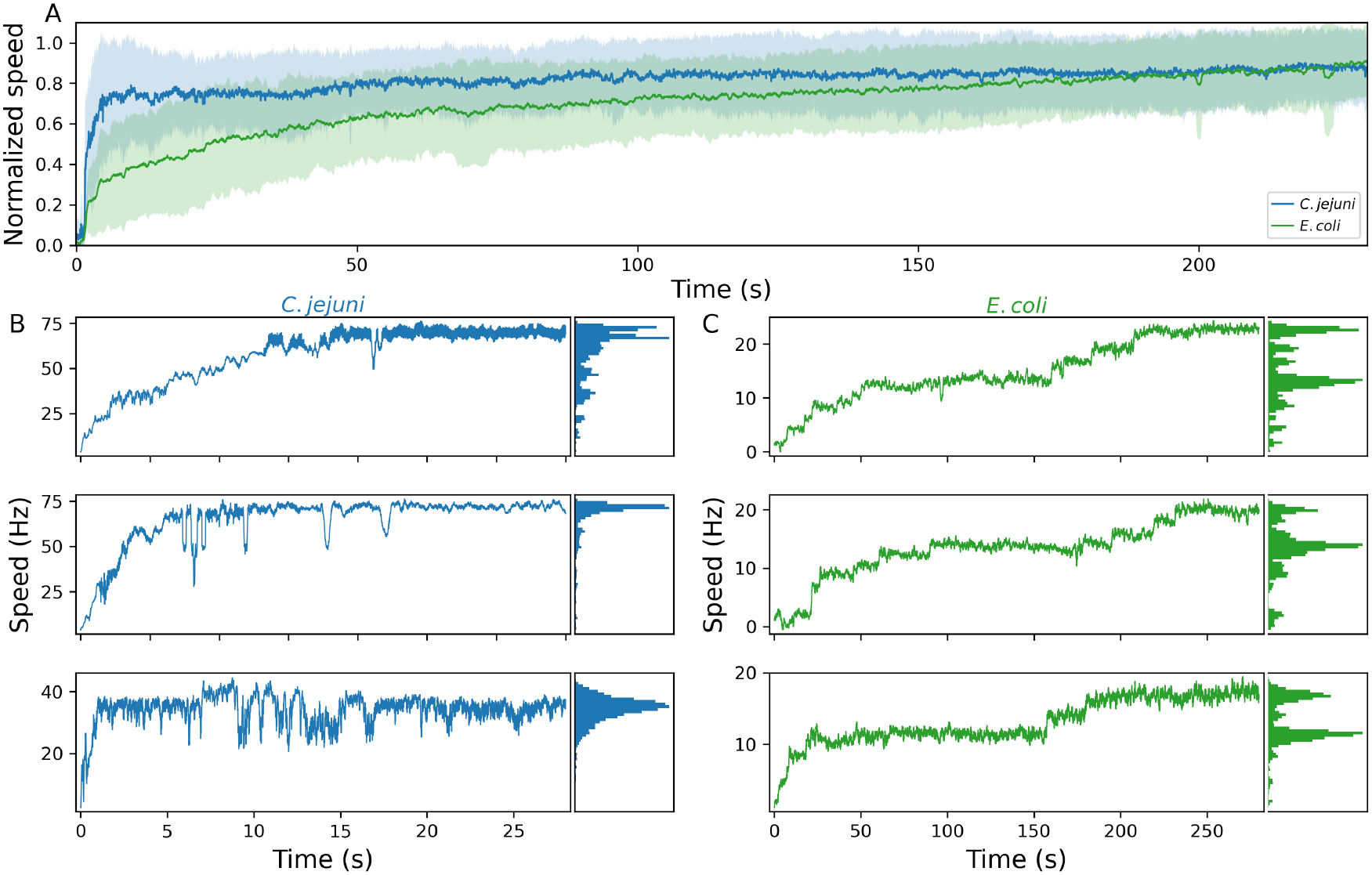
Resurrection experiments. A) Average resurrection traces for *C. jejuni* (blue) and *E. coli* (green). For each motor, angular speed was normalized by the average steady state speed measured before CCCP treatment. Resurrection traces were aligned using the detected onset of renewed rotation. Curves show the average normalized angular speed across motors after smoothing; shaded regions indicate the standard deviation. Data include 16 traces for *C. jejuni* and 18 traces for *E. coli*. *E. coli* resurrection traces were taken from ref. [30]. B, C) Examples of individual resurrection speed traces, for *C. jejuni* (B) and *E. coli* (C), shown from the detected onset of rotation, with corresponding histograms of the speeds visited during the displayed interval. Note the different time scales in B and C; recovery kinetics are compared directly in A.

Speed recovery in *C. jejuni* also exhibited less distinct steps than in *E. coli*. Plateaus between transitions were often absent, and clearly discernible steps were observed in only ∼40% of the motors (Fig. 4B,C). This limited separation of rotational states is consistent with the uncertainty in our estimates of active stator number (Fig. 3J) and suggests that the underlying activity changes are more heterogeneous or less readily resolved than stator association and dissociation in *E. coli*.

## Discussion

We developed a bead assay compatible with *Campylobacter jejuni*, enabling the first direct measurements of torque output and dynamic behavior in the flagellar motor of this organism. Our measurements reveal two main differences from the dominant picture derived from *E. coli* and *Salmonella*. First, the *C. jejuni* motor generates exceptionally high torque and exhibits a torque–speed relationship distinct from those of these model systems. Second, its torque-generating activity remains dynamic despite structural evidence for a highly occupied scaffolded stator ring. Together, these findings provide a functional counterpart to recent structural studies of the *C. jejuni* motor [20, 21, 31].

The extrapolated stall torque of the *C. jejuni* motor was approximately 3600 pN·nm, substantially greater than the values typically reported for *E. coli* and *Salmonella* and comparable to the high torque measured in the sodium-driven motor of *Vibrio alginolyticus* [11,12,32,33]. This measurement is consistent with structural models that predicted approximately three times greater torque than in enteric motors based on the larger number of stator units and the wider radius at which they act on the expanded C-ring [4,20]. However, over more than two orders of magnitude in viscous load, the torque–speed relationship differed markedly from the canonical CCW relationship of *E. coli* and the concave-down relationships reported for *Salmonella*, *V. alginolyticus*, and the wild-type dual-stator motor of *P. aeruginosa* (see Fig. S4) [11, 12, 32, 34, 35]. In the most pronounced cases, these relationships exhibit the characteristic plateau–knee profile, commonly interpreted as a crossover from a high-load, near-quasistatic regime to a low-load regime in which internal kinetic transitions limit rotation speed [13, 14]. By contrast, torque in *C. jejuni* declined steadily with increasing speed and was well approximated by a linear relationship over the measured range. Although substantial motor-to-motor variability and limited sampling at intermediate loads prevent us from excluding more subtle curvature, the data show little evidence for the pronounced concave-down curvature characteristic of these previously measured torque-speed relationships.

We propose several mechanisms that could contribute to this linear torque–speed curve. Notably, the *E. coli* motor itself exhibits markedly different torque–speed relationships in its two rotational states: the canonical CCW relationship is strongly concave-down, whereas the CW relationship is approximately linear [36]. Thus, substantially different curve shapes can arise from the same MotAB stator machinery. One possible origin is altered mechanochemical timing within the stator cycle. Models have shown that the shape of the torque–speed relationship can depend on the timing of chemical transitions relative to torque-generating motion [37, 38]; more recently, configuration-dependent ion gating has been proposed to generate a concave-down relationship by shortening torque-free waiting phases, while weaker gating yields a more nearly linear relation [39]. A different mechanical explanation attributes the directional asymmetry of *E. coli* to differences in hook compliance [40]. The linear CCW relationship observed here could therefore reflect differences in mechanochemical timing, motor–load compliance, or both, rather than requiring a fundamentally different torque-generating mechanism.

The hook-compliance mechanism is particularly relevant for *C. jejuni*. In the model of Xing et al., a soft elastic link between the rotor and the viscous load allows the motor and load to move on different timescales, producing a regime of nearly constant torque at high load [14]. Das et al. recently showed that increasing the stiffness of this linkage can collapse this timescale separation, suppress the torque plateau, and reproduce a more nearly linear torque–speed relationship [40]. The *C. jejuni* hook protein FlgE contains additional domains that form extensive intra-and inter-protofilament contacts and are predicted to increase hook stiffness and mechanical strength [41]. A stiffer *C. jejuni* hook could therefore transmit changes in rotor motion more directly to the external load, reducing the mechanical buffering that contributes to the pronounced concave-down torque–speed relationship in model systems.

Stator anchoring may provide a parallel mechanical contribution. In *E. coli*, compliance in the linkage between MotB and the peptidoglycan has been proposed to permit movement of individual stator units [14, 42]. A theoretical model incorporating compliant stator anchoring showed that soft stator springs reduce destructive interference between stator units at high speed, indicating that stator-anchoring compliance can influence collective motor output [42]. Other models have shown that the effective drag associated with stator motion can also influence how torque is transmitted to the rotor and the resulting torque–speed relationship [43]. By contrast, the stator units of *C. jejuni* are incorporated into an extensive periplasmic scaffold and enclosed by proteinaceous cage structures [4, 20, 21, 31]. This architecture could alter both the compliance and mobility of the stator units, as well as their mechanical coupling to neighboring torque-generating units. Altered stator mechanics could therefore contribute to the distinct torque–speed relationship of the *C. jejuni* motor. More generally, the torque–speed relationship of an intact motor can reflect intrinsic stator mechanochemistry, motor–load compliance, and collective stator mechanics, and the relative contributions of these factors remain unresolved [3, 43, 44].

Our time-resolved measurements revealed frequent stepwise changes in angular speed under every load tested. In *E. coli*, similar steps reflect stochastic stator unit association and dissociation, with stator occupancy regulated by mechanical load [15–18, 30]. In *C. jejuni*, however, structural reconstructions resolve 17 stator-associated positions within the scaffolded stator ring [4,20,21]. We therefore propose that the observed steps reflect transitions between functionally active and inactive states of stator units that remain physically associated with the motor. Under the assumption that each detected step corresponds to the activation or inactivation of one stator unit, we infer an average of 11 ± 4 active stators, fewer than the 17 stator-associated positions resolved structurally, consistent with some structurally associated stators not generating torque at a given moment.

The inferred number of active stator units showed no clear dependence on viscous load, in contrast to the load-dependent changes in stator occupancy observed in *E. coli*. However, substantial motor-to-motor variability and uncertainty in detecting and assigning individual steps mean that these data do not exclude mechanosensitive regulation. Our measurements compare populations of motors carrying different bead sizes, whereas controlled changes in load applied to the same motor would provide a more sensitive test [17].

The resurrection experiments provide complementary evidence that stator dynamics in *C. jejuni* differ from sequential stator recruitment in *E. coli*. Following PMF restoration, the *C. jejuni* motor reached 50% of its pre-treatment speed in approximately 3 s, compared with approximately 25 s in *E. coli*. Recovery in *C. jejuni* also often lacked the well-separated plateaus characteristic of sequential stator unit incorporation in *E. coli*. These observations are consistent with rapid reactivation of pre-associated stator units rather than their slower recruitment from a membrane pool.

Several mechanisms could allow a structurally assembled stator ring to dynamically regulate torque. The MotB plug domain might reversibly gate ion conduction without requiring stator dissociation, although its established role is principally to prevent ion leakage before the assembly of the stator unit [45, 46]. Alternatively, conformational changes within the stator or at the stator–rotor interface could transiently disengage torque generation while leaving the complex incorporated within the scaffold, reminiscent of motor-associated brakes and clutches that regulate torque without stator loss in other species [47–50]. The extensive connections between stator units and their surrounding scaffold may also provide sites through which stator activity could be modulated without physical exchange [20, 21, 31]. Such regulation could allow the motor to maintain a reserve of inactive stator units that can be mobilized rapidly without requiring new stator assembly. Auxiliary viscosity-responsive regulators may also contribute: VidB has been proposed to act as a brake or clutch at low viscosity, while VidA forms part of a motor-associated V-ring implicated in viscosity sensing [51, 52]. Whether these regulators control the active stator states inferred here, or the conditions under which this reserve capacity is engaged, remains unknown.

Unlike many bacteria, which slow as external viscosity increases, *C. jejuni* can maintain or increase its swimming speed in viscous environments [51, 53, 54]. The high torque measured here likely contributes by allowing the motor to sustain rotation against large mechanical loads, but efficient motility also depends on adaptations at the whole-cell level. The helical cell body counter-rotates relative to the flagella and acts as an additional screw-like propeller, while wrapped-mode swimming can align the thrust produced by the two polar flagella [55]. The large basal disk may further support motility by stabilizing the motor during unwrapping of the leading flagellar filament from the cell body during directional reversals [56]. High motor torque may therefore act in concert with these whole-cell mechanical adaptations to support efficient motility in mucus and other viscous environments.

More broadly, our findings illustrate the diversity of evolutionary strategies available for adapting bacterial flagellar motors to different mechanical and ecological demands. In *E. coli* and other well-studied motors, stator exchange enables adaptation through dynamic remodeling of motor composition [15–17, 30]. In *C. jejuni*, by contrast, a high-capacity stator ring appears to be assembled and structurally stabilized in advance, with our measurements consistent with regulation through reversible changes in stator activity. These results establish that pronounced concave-down torque–speed behavior and load-dependent regulation through stator exchange are not universal features of bacterial flagellar motors, and reinforce the view of the flagellar motor as a modular and evolvable machine whose mechanical behavior has diversified alongside its structure [5, 6].

## Materials and Methods

### Strain Construction and Genetic Modifications

The strain used in this study was constructed as described previously [55] and used in subsequent work [20]. The parent strain was DRH212, a streptomycin-resistant derivative of *C. jejuni* 81-176 [55,57]. The genotype of the strain relative to DRH212 was Δ*flaA flaB^S397C^ flhF^D321A^* Δ*cheY*. Deletion of *flaA* reduced the length of the flagellar filament to 200-1000 nm, corresponding to the length of FlaB [55]. The S397C substitution in *flaB* enabled maleimide coupling, as described below. The D321A substitution in *flhF* increases the frequency of flagellar motors at nonpolar positions, facilitating bead measurements away from the cell–surface interface [20, 58]. Deletion of *cheY* produced motors that rotated exclusively in the counterclockwise direction.

### Bacterial culture

*C. jejuni* was cultured on Mueller-Hinton (MH) agar plates (1.4% agar) supplemented with 10 µg/mL trimethoprim (Sigma-Aldrich, T7883-5G), following a previously published protocol [59]. Trimethoprim stock solutions were prepared at 10 mg/mL in dimethyl sulfoxide and stored at 4°C until use. A frozen stock was prepared from a swab in charcoal transport agar: the swab was streaked onto an MH plate and incubated overnight. Bacteria were then harvested from the plate and stored in cryovials containing 80% MH broth (BD Difco, 275730) and 20% glycerol (Sigma-Aldrich, G9012-1L). This frozen stock was used to seed all plates for bead assay experiments.

For each bead assay experiment, a small amount of frozen stock was scraped from the cryovial using a sterile pipette tip or toothpick and spread over a small area of an MH agar plate. The plate was incubated in a sealed pouch with a gas-generating sachet (BD GasPak EZ CampyPouch System, 260685) and a moist paper towel at 37°C for 16-48 hours. The resulting bacterial patch was collected with an inoculation loop and restreaked onto a fresh MH plate, which was incubated overnight under the same conditions. If needed, a third plate was prepared by streaking from the second. Additional passages were performed at most twice following the first full plate.

### Sample Preparation

After growth, bacteria were rinsed from the MH agar plate using MH broth. The resulting suspension was centrifuged at 8000 rpm for 2 min, and the pellet was resuspended in 500 µL phosphate-buffered saline (PBS; Sigma-Aldrich, 79383-250ML). This washing step was repeated once more, and the suspension was adjusted to an optical density (OD_600_) of 2. To biotinylate the truncated flagellar filaments, 25 µL of 20 mM maleimide-PEG2-biotin linker (ThermoFisher Scientific, EZ-Link Maleimide-PEG2-Biotin, 21901BID) in PBS was added to the bacterial suspension and incubated for 10 min to allow coupling to cysteine residues [22, 55]. Bacteria were then washed twice by centrifugation (8000 rpm, 2 min) and resuspended in 500 µL PBS.

Tunnel slides were prepared in advance following a previously described protocol [22]. Each tunnel slide was assembled from two glass coverslips (Menzel-Gläser 631-0851) separated by a parafilm spacer. The top coverslip contained two laser-cut holes (2 mm diameter) to allow flow into and out of the tunnel. PDMS blocks with 2 mm holes were plasma bonded over the holes to support Luer fittings, which connected the slide to a peristaltic pump for flow control. The tunnel slide was assembled by lightly melting the parafilm spacer between the holed coverslip and a second coverslip. Luer fittings were then inserted into the PDMS blocks.

To clean and prepare the tunnel slide, 1 mL of 70% ethanol followed by 1 mL of MilliQ water was flushed through the chamber. Next, 250 µL of Poly-L-lysine (PLL; Sigma-Aldrich, P4707-50ML) was added and incubated for 5 min to coat the lower surface. The slide was washed with 500 µL PBS, followed by 200 µL of the bacterial suspension, which was allowed to settle for approximately 10 min. To maximize surface contact between the bacteria and glass, the tunnel was washed rapidly with 2 mL PBS to force the bacteria down. Next, 3 mL of MH broth was flushed through to remove loosely attached cells.

To label the biotinylated flagella, 200 µL of streptavidin beads in MH broth were added to the chamber and incubated for 5–10 min. Bead sizes used included: 250 nm and 450 nm (Ademtech, 03131 and 03150), 800 nm (Abvigen, ABPS-0080-SA), 1100 nm (Thermo Fisher Scientific, 65001), 1400 nm (Sigma-Aldrich, 49532-5ML), and 2700 nm (Thermo Fisher Scientific, 65305). The slide was then gently washed with 1 mL MH broth, and the chamber was left open to allow passive exchange of medium. After each acquisition (typically 5 min in duration), 250 µL of fresh MH broth was added to the tunnel to maintain oxygenation in the flow cell.

### Data Collection

Bead rotation was characterized using a custom-built bright-field microscope. Illumination was provided by a light-emitting diode (Thorlabs M660L4) focused onto the sample, and transmitted light was collected using a 100× oil-immersion objective (NA = 1.46, Zeiss 420792-9800-720). The resulting image was relayed to a high-speed CMOS camera (Optronis CL600X2-FULL-M-FM), capturing defocused bead images suitable for rotation tracking. Videos of bead rotation were acquired at frame rates between 1000 and 5000 frames per second (FPS) during steady-state rotation.

To collect resurrection data, motors were labeled with 1400 nm beads, and 5 µM carbonyl cyanide m-chlorophenyl hydrazone (CCCP) in MH broth was added to the flow cell to dissipate the PMF. After incubation for 5–8 minutes, the slide was flushed with fresh MH broth. Bead rotation resumed, and the resurrection process was recorded at 2000–8000 FPS for a duration of 5 minutes.

### Data Analysis

Motor angular speed and torque were extracted from video recordings of bead rotation [22, 60]. Beads were tracked to yield time series of *x* and *y* positions for the bead center. Lateral drift was corrected by fitting a spline to the mean bead position within sequential time windows (typically 30 windows over a 5-minute acquisition) and subtracting the shift in mean position relative to the first window. The resulting trajectory was then circularized to correct for ellipticity caused by out-of-plane bead motion. The short axis of the ellipse was stretched to match the long axis, and angular position was calculated as *θ* = atan2(*y, x*), with the trajectory centered at (0, 0). The angle was unwrapped to produce a continuous angular trajectory, and the instantaneous angular velocity at time point *i* was calculated as Ω*_i_* = (*θ_i_*_+1_ − *θ_i_*)*/*Δ*t*, where Δ*t* is the time interval between frames. Motor torque *τ_i_* was computed as *τ_i_* = (*γ_b_* +*γ_f_* +*γ_r_*) Ω*_i_*, where *γ_b_* is the viscous drag on the bead, *γ_f_* is the drag from the filament stub, and *γ_r_* accounts for internal friction within the rotor. Angular speeds are reported in Hz throughout the figures and text, corresponding to *f_i_* = Ω*_i_/*2*π*, while torque was calculated using angular velocity in rad/s.

The bead drag coefficient *γ_b_* was calculated as in [32]:

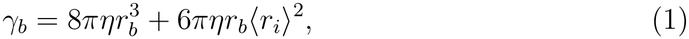

where *η* is the dynamic viscosity of the medium (approximated as that of water 1.0 · 10*^−^*^3^ Pa s), *r_b_* is the bead radius, and ⟨*r_i_*⟩ is the mean radius of bead rotation. Bead sizes were independently measured by transmission electron microscopy; Fig. S1 shows boxplots of bead diameters and representative images, and Table S2 lists average values of *γ_b_* for each bead size. Unlike previous work in *E. coli* [17, 60], we omitted corrections to *γ_b_* for proximity to the cell surface [61, 62], as this distance was unknown in our setup.

The filament drag coefficient *γ_f_* was calculated using the expression from [32]:

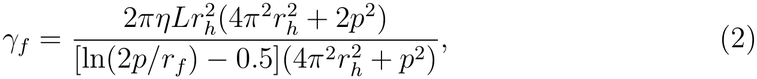

where *L* is the contour length of the filament (0.7 µm in the Δ*flaA* mutant [55]), *r_f_* is the filament radius (9 nm from cryo-EM [63]), *r_h_* is the helical radius, and *p* is the pitch. Assuming one full helical turn over the 2.5 µm wild-type contour length [55,64] and a helix angle of 44*^◦^* [55], we estimate *r_h_* = 0.27 µm and *p* = 1.8 µm, yielding *γ_f_* = 0.09 pN nm s/rad.

The rotor drag coefficient *γ_r_*, which reflects friction internal to the motor, was estimated by scaling from the *E. coli* value of 0.10±0.01 pN nm s/rad [65,66]. Given that the rotor of *C. jejuni* is approximately 10 nm larger in diameter than that of *E. coli*, and that drag on a rotating cylinder scales with diameter squared [67], we approximated *γ_r_* in *C. jejuni* as 0.15 pN nm s/rad. Unless otherwise specified, angular speed and torque traces plotted outside the resurrection analysis were smoothed using a Savitzky–Golay filter with a 0.5 s window.

For resurrection analysis, angular speed traces were normalized by the average steady-state angular speed measured before CCCP treatment. *E. coli* resurrection traces were taken from ref. [30] and treated as previously processed speed traces. Raw *C. jejuni* resurrection traces were smoothed using a 0.05 s running window to generate comparable processed speed traces. These processed traces were used for onset detection, averaging, representative traces, and histograms. The onset of renewed rotation was defined as the first sustained interval after the recorded resurrection start point in which speed exceeded the larger of 1 Hz or 5% of the post-start 95th percentile speed for at least 0.75 s. For the average traces in Fig. 4A, individual traces were aligned to this detected onset and plotted beginning 1.5 s before onset. *C. jejuni* traces acquired at 8000 FPS were downsampled to an effective 2000 FPS before averaging. Representative traces and histograms in Fig. 4B,C are shown from the detected onset of rotation.

### Detection and Quantification of Discrete Motor Speed Steps

We processed angular speed traces to detect discrete transitions in motor speed. To reduce high-frequency noise while preserving step-like changes, angular speed data were smoothed using a Savitzky-Golay filter (second-order polynomial, 0.25 s window). Discrete transitions were then detected using the Python package ruptures [68], employing the PELT algorithm with an L2 cost model. The penalty parameter was adjusted for each experimental condition to account for differences in signal length and noise level. Candidate transitions were further filtered by retaining only those with magnitude greater than *ω*_max_*/*34, where *ω*_max_ is the maximum filtered angular speed in the trace. Between each pair of retained transitions, the angular speed was replaced by its median value, producing a piecewise-constant angular speed trace *ω*(*t*). The single-stator speed increment, Δ*ω*, was estimated as the median absolute change in *ω*(*t*) between adjacent levels.

## Acknowledgements

We thank J. Finkel for collecting the transmission electron microscopy data. W.H.H, E.J.C., M.B, and F.P. were supported by Human Frontier Science Program grant RGP0028/2021-HOCHBERG. E.J.C. and M.B. were further supported by UK Medical Research Council grant MR/V000799/1, and F.P. was further supported by the ANR BaElMec ANR-23-CE30-0010 project grant. S.L. was supported by the UChicago-CNRS Joint PhD Research Program, which is a part of the IRC Discovery. A.L.N. and A.G. were supported by the ANR PHYBABIFO project grant ANR-22-CE30-0034. A.L.N was further supported by the ANR PHYBION project grant ANR-23-ERCB-0005-01 and the Impulscience project PHYBION of the Bettencourt Schueller Foundation. The CBS is a member of the France-BioImaging (FBI) and the French Infrastructure for Integrated Structural Biology (FRISBI), two national infrastructures supported by the French National Research Agency (ANR-10-INBS-04-01 and ANR-10-INBS-05, respectively).

