## Supplementary Information for "Torque generation and regulation in the flagellar motor of *Campylobacter jejuni* differ from those of model organisms"

#### S1 Bead sizes

We used transmission electron microscopy (TEM) to verify the size distributions of the beads used in this study. For each bead size, we measured the diameters of at least 100 individual beads. Figure S1 shows box plots summarizing these measurements alongside representative TEM images for each bead type.

#### S2 Torque-speed plots by bead size with individual motor measurements

Fig. S2 shows torque-speed plots for each bead size, with individual motor measurements represented as colored points. For each bead size, a linear regression constrained through the origin was fitted to torque as a function of rotational frequency. Because frequency is plotted in Hz, the fitted slope corresponds to $2\pi\gamma$; division by $2\pi$ yields the drag coefficient $\gamma$ in pN·nm·s/rad. Because torque was calculated using the theoretical drag assigned to each motor, these load-line values provide a summary of the effective drag within each bead-size group rather than an independent drag measurement. These regressions provide bead-size-specific estimates of drag, shown in Table S1, along with the average angular speed and torque for each condition. Fig. S3 presents

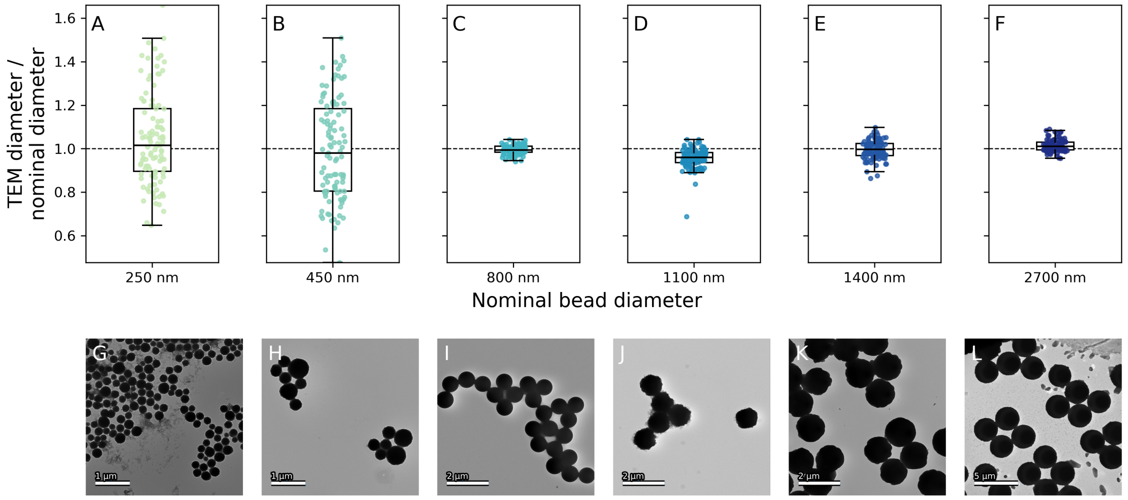


Figure S1: Transmission electron microscopy (TEM) analysis of bead sizes. A–F) Box plots showing TEM-measured bead diameters normalized by the nominal bead diameter for each bead size; the dashed line indicates the nominal diameter. The box indicates the interquartile range and the whiskers are drawn to the farthest data point within 1.5 × the interquartile range. G–L) Representative TEM images corresponding to each bead size shown in panels A–F.

the full torque–speed curve for *C. jejuni*, combining data from all motors characterized.

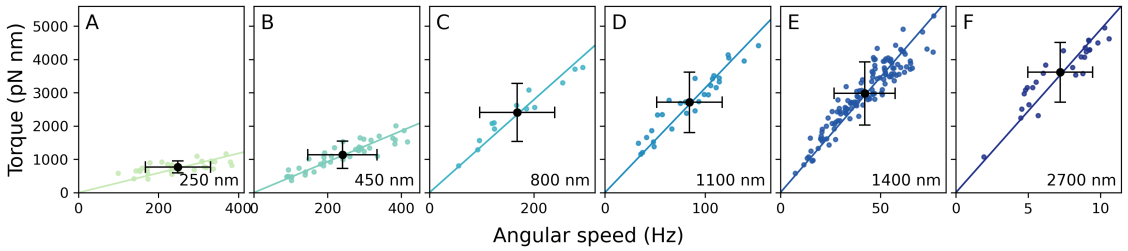


Figure S2: Torque-speed plots for each bead size, as indicated by the panel label. A–F) Colored points show average angular speeds and torques for individual motors. The solid line is a linear regression (forced through the origin) representing the average load line for each dataset. Black points indicate the average angular speed and torque across motors for that bead size; error bars show standard deviations.

### S3 *C. jejuni* torque–speed curve compared to other bacterial species

Fig. S4 compares the torque–speed curve of *C. jejuni* with those of several other bacterial species. These include both proton-driven and sodium-driven motors from Gram-negative and Gram-positive organisms. Each point represents the average motor torque and speed at a given load (defined by bead size and/or medium viscosity).

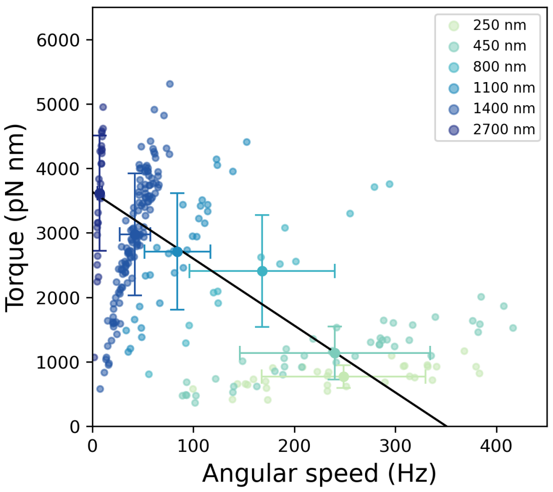


Figure S3: Torque-speed curve of *C. jejuni* incorporating all motors characterized in this study. Each small colored point represents the torque and angular speed of an individual motor, color-coded by bead size. Large colored points show the average torque and angular speed for each bead size, with standard deviations as error bars. The black line is the linear regression fitted to the bead-size averages, as shown in Fig. 2C.

| Bead Size (nm) | 250 | 450 | 800 | 1100 | 1400 | 2700 |
|---|---|---|---|---|---|---|
| TEM diameter (nm) | 262 ± 53 | 450 ± 100 | 797 ± 17 | 1053 ± 50 | 1390 ± 58 | 2740 ± 73 |
| Fit-based drag (pN nm s/rad) | 0.461 ± 0.021 | 0.720 ± 0.023 | 2.128 ± 0.064 | 4.82 ± 0.13 | 10.54 ± 0.13 | 74.2 ± 2.0 |
| Theoretical drag (pN nm s/rad) | 0.52 ± 0.15 | 0.77 ± 0.14 | 2.25 ± 0.24 | 5.14 ± 0.83 | 11.1 ± 1.5 | 78 ± 10 |
| Angular Speed (Hz) | 252 ± 78 | 235 ± 94 | 168 ± 72 | 82 ± 36 | 44 ± 14 | 7.4 ± 2.3 |
| Torque (pN nm) | 770 ± 170 | 1090 ± 390 | 2310 ± 830 | 2560 ± 970 | 3010 ± 760 | 3520 ± 860 |

Table S1: Average bead diameter, drag, angular speed, and torque for each bead size. Values in the row labeled "TEM diameter" are the mean ± standard deviation of bead diameters measured by TEM. Values in the row labeled "Fit-based drag" were estimated from the slope of the linear regressions in Fig. S2; uncertainties represent the standard error of the slope. Values in the row labeled "Theoretical drag" were calculated using the theoretical drag of Eq. 1, based on the TEM-measured bead radius and the individual trajectory radius of each motor; uncertainties represent the standard deviation across motors. Note: The drag values labeled "Theoretical drag" were used in all torque and torque–speed calculations presented in the main text.

### S4 *C. jejuni* resurrection traces

Fig. S5 shows the resurrection traces of *C. jejuni* that make up the average resurrection trace plotted in Fig. 4D. The left column shows the whole traces starting from where there is an increase from zero angular speed. The middle column shows the corresponding portions of the data in the left column where the greatest change in angular speed is observed. The right column shows the resurrection traces in the middle column plotted as a histogram of angular speeds. These histograms highlight the presence of dwell speeds in some of the resurrections. The traces marked with an asterisk are those plotted in Fig. 4A–C.

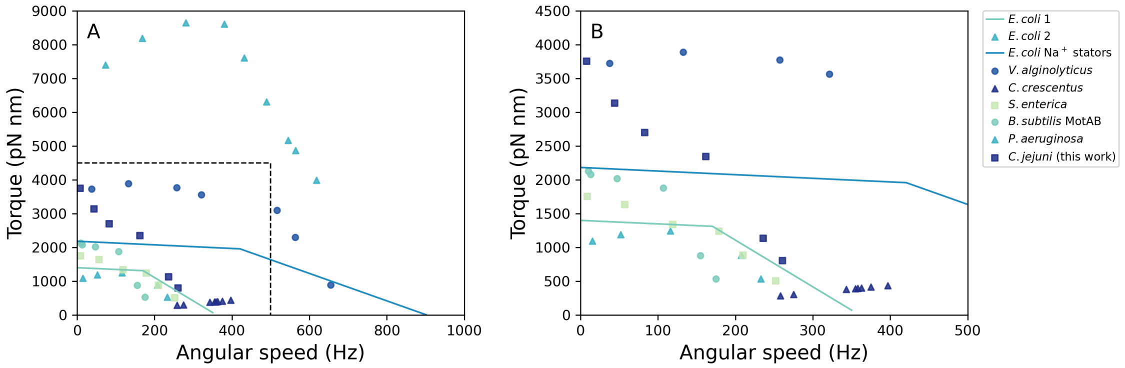


Figure S4: Comparison of torque–speed curves across bacterial species. A) Curves shown include *Escherichia coli* from two studies [?, 1], *E. coli* expressing chimeric sodium-driven stators [2], *Vibrio alginolyticus* [?], *Caulobacter crescentus* [3], *Salmonella enterica* [?], the MotAB stator of *Bacillus subtilis* [?], *Pseudomonas aeruginosa* [?], and *C. jejuni* (this work). Each point corresponds to an average torque and angular speed for a given drag condition. The continuous curves are fits to individual motor data. The dashed box marks the region enlarged in B. B) Zoomed-in view of the low-speed, low-torque region shown in panel A.

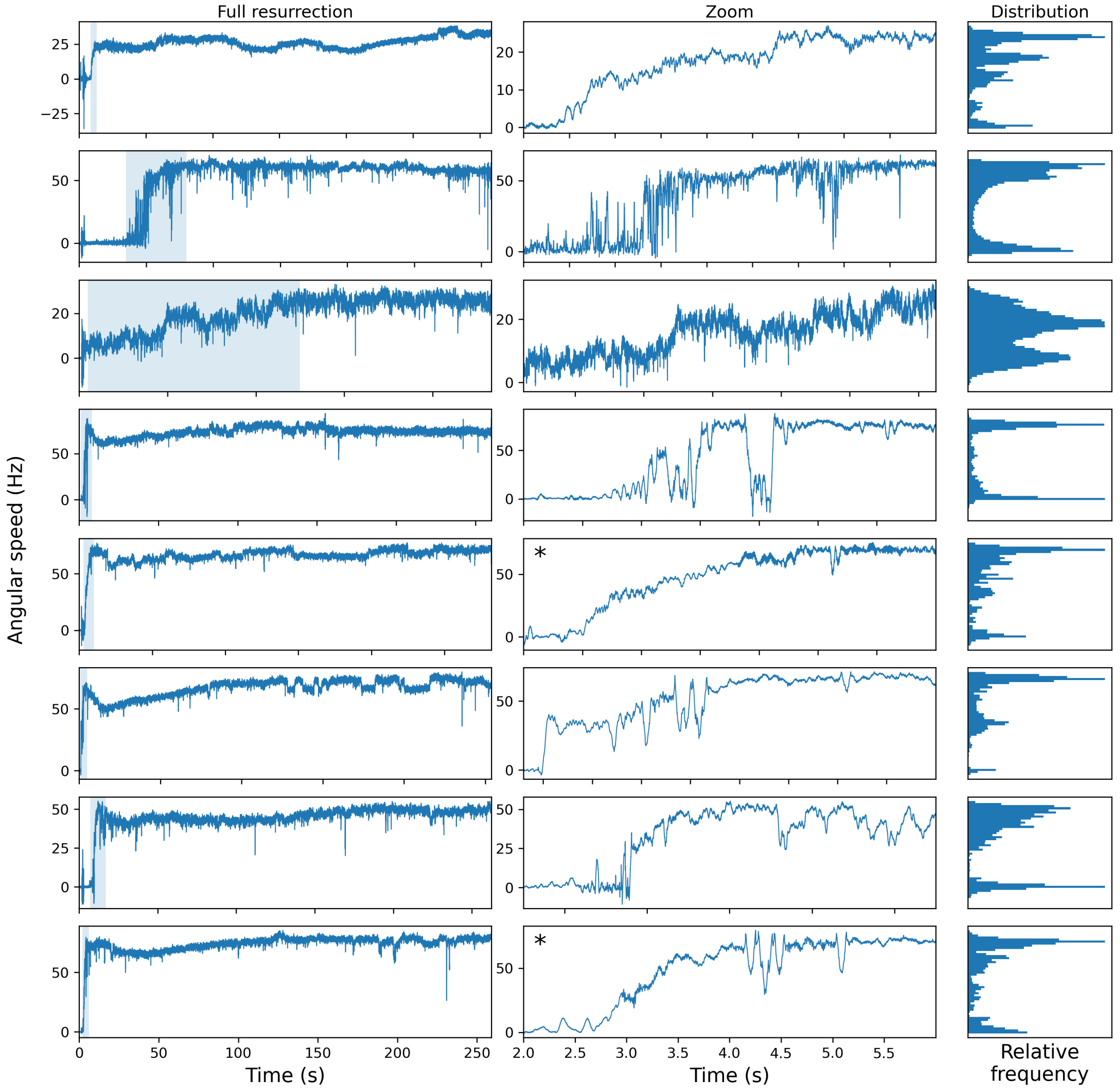


Figure S5: Continued below.

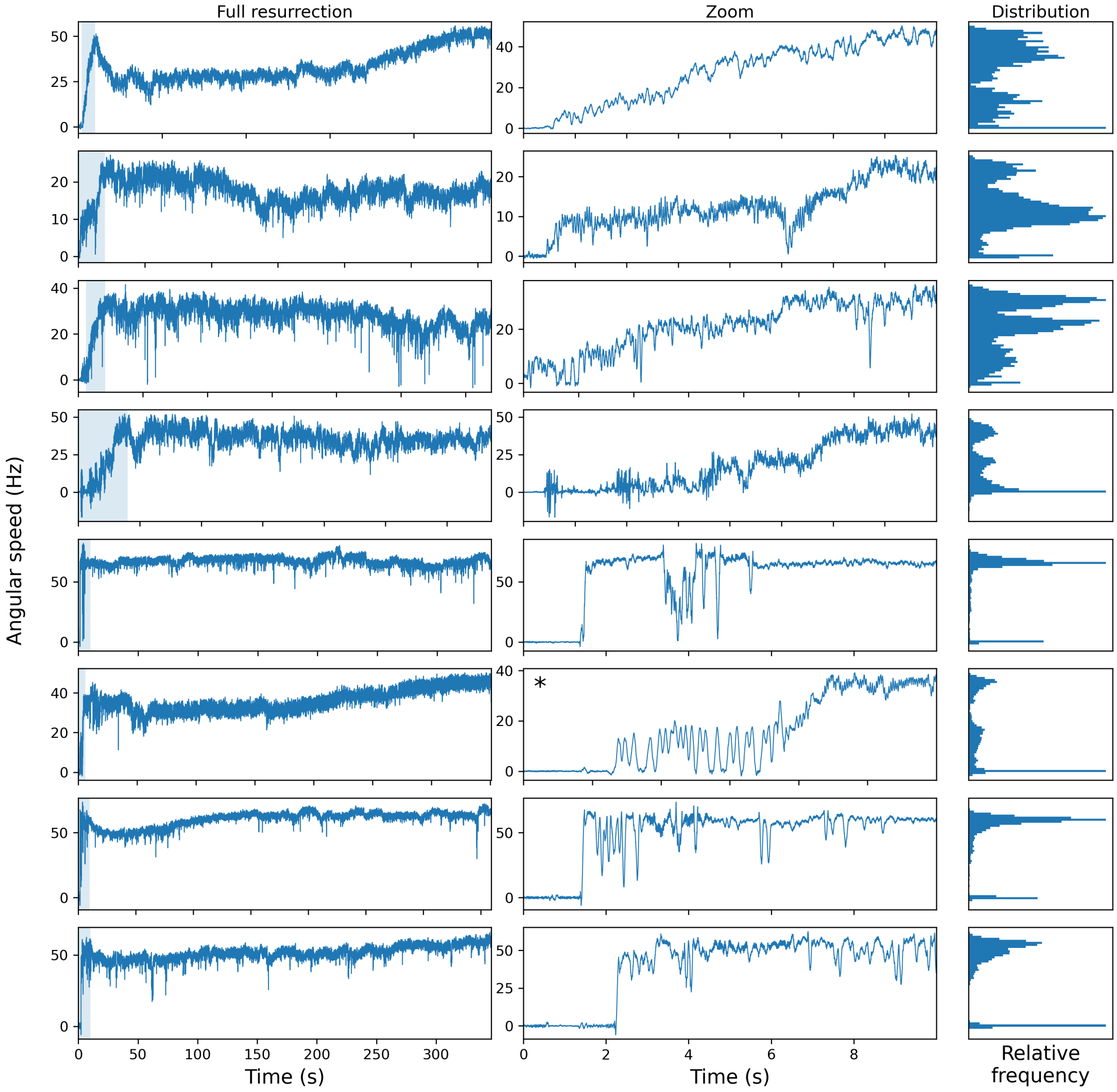


Figure S5: (continued) Individual resurrection traces of *C. jejuni*. Left column: Full resurrection traces. Middle column: Portions of resurrection traces in the left column where there is the greatest change in angular speed. The light blue shaded region in the left column is the portion of the resurrection trace shown in the middle column. Right column: Angular speed histograms of the corresponding angular speed traces in the middle column. A Savitzky-Golay filter with a 0.1 s window and a polynomial order of 3 was used. The starred traces are the traces in Figure 4A–C.

### References


[1] Ekaterina Krasnopeeva, Lucas Le Nagard, Uriel E. Barboza-Perez, Wilson Poon, Chien-Jung Lo, and Teuta Pilizota. Nonlinear dependency of the bacterial flagellar motor speed on proton motive force and its consequences for swimming. *bioRxiv*, 2026. Version 2, posted 2 February 2026.

[2] Yuichi Inoue, Chien-Jung Lo, Hajime Fukuoka, Hiroto Takahashi, Yoshiyuki Sowa, Teuta Pilizota, George H. Wadhams, Michio Homma, Richard M. Berry, and Akihiko Ishijima. Torque-speed relationships of $Na^{+}$-driven chimeric flagellar motors in Escherichia coli. *Journal of Molecular Biology*, 376(5):1251–1259, 2008.

[3] Guanglai Li and Jay X. Tang. Low flagellar motor torque and high swimming efficiency of *Caulobacter crescentus* swarmer cells. *Biophysical Journal*, 91(7):2726–2734, 2006.
